# fMRI replicability depends upon sufficient individual-level data

**DOI:** 10.1101/352633

**Authors:** Derek Evan Nee

## Abstract

The replicability of findings drawn from functional magnetic resonance imaging (fMRI) data have increasingly been called into question. Concerns have been raised that historically, sample sizes have been insufficient to produce adequate power, leading to unreliable results. Recently, Turner and colleagues demonstrated that even with large sample sizes, the replicability of fMRI can be worryingly low. However, their datasets featured low amounts of data at the individual-level. Here, I demonstrate that replicability depends critically on sufficient individual-level sampling. I show that fMRI can have strong replicability even at modest sample sizes when individuals are adequately sampled, but that inadequate individual-level sampling leads to poor replicability. These data indicate that fMRI replicability cannot be judged solely on sample size, and that adequate sampling at the individual-level is a critical design consideration.

---

The reproducibility of task-based functional magnetic resonance imaging (fMRI), or lack thereof, has become a topic of intense scrutiny^1,2^. Relative to other human techniques, fMRI has high costs associated with data collection, storage, and processing. To justify these costs, the inferences gained from fMRI need to be robust and meaningful. Hence, although large, sufficiently powered datasets may be costly, this is favorable to collecting many insufficiently powered datasets from which reliable conclusions cannot be drawn. However, it can be difficult to determine *a priori* how much data are needed. Although power analyses can help^3^, accurately calculating power itself requires an appropriate estimate of the expected effect size, which can be hard to obtain if previous studies had insufficient data to produce reliable effect size estimates. Furthermore, mechanistic basic science explores novel phenomena with innovative paradigms such that extrapolation of effect sizes from existing data may not be appropriate.

In light of these issues, many studies rely on rules-of-thumb to determine the amount of data to be collected. For example, Thirion et al^4^ suggested that twenty or more participants are required for reliable task-based fMRI inferences. Turner et al^5^ recently pointed out that such recommendations are outdated, and set out to empirically estimate replicability using large datasets. The authors found that even datasets with one-hundred or more participants can produce results that do not replicate, suggesting that large sample sizes are necessary for task-based fMRI.

It is typical for considerations of power in task-based fMRI to focus on sample size. This is because between-subject variability tends to dominate within-subject variability, such that sampling more subjects is often a more effective use of time than scanning individuals for longer^3,4^. Large task-based fMRI data collections such as the Human Connectome Project (HCP) have used batteries of tasks wherein each task is scanned on the order of ten minutes^6^. Such batteries operate under the assumption that within-subject variability, which diminishes with scan time, can reach appropriately low levels within a relative short period. However, using data from the HCP and other data of similar durations, Turner et al^5^ demonstrated that task-based fMRI can be unreliable.

With the rising popularity of resting-state fMRI, investigators have examined the duration of resting-state data needed for reliable parameter estimates. Some have suggested that parameter estimates are stable after 5-10 minutes of resting-state scans^7^, although more recent data suggest 30-40 minutes are needed^8,9^. In either case, parameters estimated from rest use the entire (cleaned) data time-series, while task-based fMRI splits the time-series into composite mental events. For example, in a rapid event-related design, there may be approximately 4-6 seconds of *peak signal* attributable to a given transient event-of-interest (e.g. a choice reaction). If twenty such events exist in a ten-minute task run, that amounts to less than two minutes of signal attributable to that task event. Although it is difficult to extrapolate from rest to task given the numerous differences between the methods, it is likely that parameter estimates in such short tasks would benefit from additional measurements at the individual-level.

To examine the impact of individual-level measurements on task-based fMRI replicability, I re-analyzed data from a recently published pair of datasets^10,11^. Each dataset estimated five contrasts-of-interest spanning main effects and an interaction in a 2×2×2 factorial design. The resultant contrasts variously load on often-studied constructs of working memory, task-switching, language, and spatial attention. These constructs have a high degree of overlap with those examined by Turner et al^5^. Previously, I suggested the reproducibility in these data were good^10,11^, but given the observations of Turner et al^5^, the sample sizes employed (n=24) should produce low replicability. On the other hand, ∼one-two hours of task data were collected for each individual, which could have facilitated reliability. To formally examine this matter, I computed the replicability measures of Turner et al^5^ on randomly sub-sampled independent datasets for the five contrasts-of-interest. I varied the amount of individual-level data from ∼ten minutes (one task run) to ∼one hour (six task runs). I also varied the sample size from sixteen to twenty-three individuals with sixteen matching the minimum examined by Turner et al^5^ and twenty-three being the maximum that can be split into independent groups in the forty-six participants examined. All data and code are available at https://osf.io/b7y9n.

Figure 1 shows the results at n=16. When only one run is included for each individual, the replicability estimates fall in the ranges reported by Turner et al^5^. However, reproducibility markedly improved with more data at the individual-level. While there are some indications of diminishing returns after four runs, there were clear benefits to more scans at the individual-level. Figure 2 reports the results at n=23, which again show clear benefits to reproducibility with more than one run. For example, the mean peak replicability with two runs (∼65%) matches observations in Turner et al^5^ at n=64. Furthermore, no contrast in Turner et al^5^ approached perfect replicability with any combination of measure, sample size, and threshold, whereas multiple combinations produced near perfect replicability for the Contextual Control contrast with as little as six runs at n=16 (Supplemental Figure 1). In the most striking such case, I find nearly 90% of the peaks replicate on average with four runs at n=23 (Supplemental Figure 2), which again exceeded the observations of Turner et al^5^ even at the largest sample size (n=121). While the differences in tasks employed here and those in Turner et al^5^ qualify direct comparisons, the data here paint a much more reliable picture of task-based fMRI at modest sample sizes when individuals are adequately sampled.

**Figure 1.**
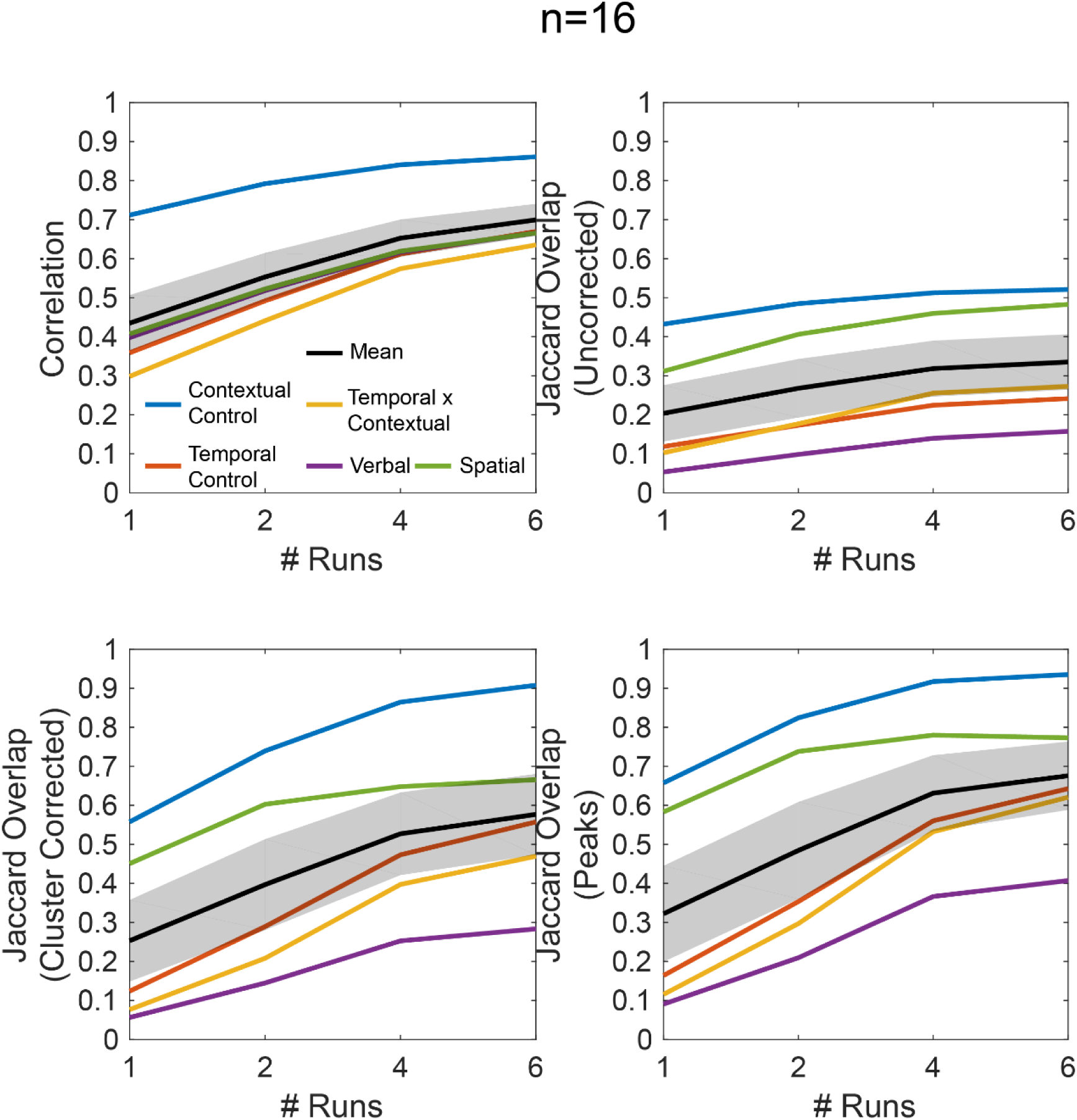
Replicability estimates at n=16. Metrics correspond to those used in Turner et al^5^. Jaccard Overlaps were calculated using conservative thresholds comparable to those reported in Turner et al^5^.

**Figure 2.**
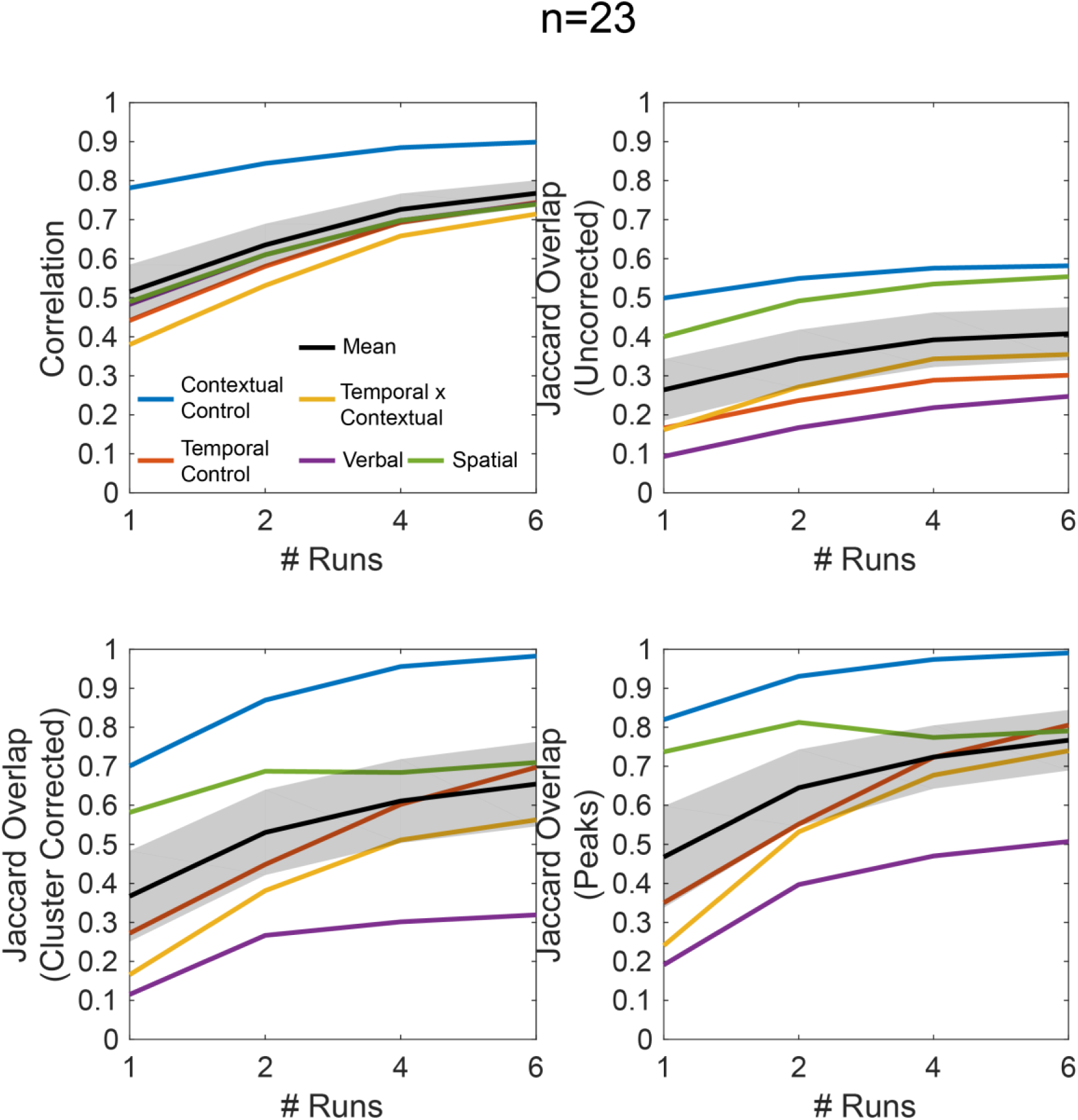
Replicability estimates at n=23. Other details match Figure 1.

These observations raise the question of how much individual-level data are needed. This is not straightforward to determine *a priori* and hinges on the ratio of within-to between-subject variability and effect magnitude (see ^12^ for demonstrations of how these factors trade-off). Concrete recommendations are rendered difficult given that these factors will vary considerably based on experimental design (including how the data are modeled), brain region, population, scanner, and scanning parameters. In the data explored here, at n=23 with six runs, peaks from the Contextual Control contrast were nearly perfectly reliable, while only half of the peaks from the Verbal contrast replicated despite these contrasts being matched for time and number of trials, demonstrating that one size does not fit all. In general, more data at the individual level are beneficial when within-subject variability is high, and between-subject variability is low^12^. Furthermore, across all of the contrasts, I observed diminishing returns after approximately four task runs, which may owe to the duration of time participants can remain attentive and still (i.e. ∼forty-minutes) and/or the point at which the within-subject variability is sufficiently low relative to the between-subject variability. Hence, forty-minutes of task may be a reasonable starting point for pilot data, from which the appropriate parameters can be estimated and used to determine proper levels of *n* and scan time.

A final question is the extent to which researchers are scanning sufficiently at the individual-level. An assay of recent studies of basic mechanistic research indicates that modest sample sizes are the norm (mean N=31.7), but few studies employ less than ten-minute scanning durations (Supplemental Figure 3). The average per task scanning duration was ∼forty-minutes, which matches the point of diminishing returns observed here. Hence, the observations of Turner et al^5^ based on short scans cannot be broadly generalized to basic science research that tends to scan much longer. However, those studies employing batteries of short tasks would do well to consider the observations of Turner et al^5^ and here, and collect more individual-level data to foster reproducibility.

## Methods

Full details of the participants, task, preprocessing, and modeling can be found in my previous reports^10,11^. Briefly, the task manipulated two forms of cognitive control (contextual control, temporal control) and stimulus domain (verbal, spatial) in a 2−2−2 factorial design. Five contrasts from the factorial design were included in this report: contextual control, temporal control, temporal control x contextual control, verbal (> spatial), and spatial (> verbal). On each block, participants performed a sequence-matching task in a given stimulus domain. Then, sub-task phases orthogonally manipulated the cognitive control demands. In the original report, we examined stimulus domain (verbal>spatial, spatial>verbal) across all trials. But here, I use only the sub-task phases so that all contrasts have the same amount of data at the individual level. A separate contrast estimate was created for each individual and each run. I included data from 46 participants, excluding participants in the original reports that did not complete all of the task runs. 23 participants performed 12 scanning runs and 23 participants performed 6 scanning runs, wherein each scanning run took approximately 10 minutes to complete. Data and code are available at https://osf.io/b7y9n.

Following the procedures of Turner et al^5^, replicability was determined by pairwise comparison of group-level t-statistic maps. For each analysis, the data were randomly split into two independent groups 500 times. Analyses varied the number of runs included at the individual level (1, 2, 4 or 6) by randomly selecting a subset of the data, and also the number of individuals (16 or 23). Extra-cranial voxels were masked out and voxels for which t-statistics could not be computed (i.e. due to insufficient signal across participants) were discarded prior to computations of replicability.

The first analysis examined the voxel-wise correlation of t-statistics across all voxels. Subsequent analyses examined Jaccard overlap on thresholded t-statistic maps where the Jaccard overlap indicates the proportion of results that replicate. Although Turner et al^5^ utilized both positive and negative activations for their Jaccard overlap calculations, here I use only positive activations given that two of the contrasts are the inverses of one another. Following Turner et al^5^, Jaccard overlap was computed at the voxel-level by first thresholding the complete group dataset and determining the number of significant voxels, *v*, at a voxel-wise threshold. This map represented the “ground truth.” Then, in each pair of sub-sampled datasets, the conjunction of the top *v* voxels was divided by their union to determine the proportion of replicated voxels.

The voxel-level procedure does not attempt to control false-positives for each group analysis. Therefore, low replicability in this measure might be anticipated by the inclusion of false-positives. So, Turner et al^5^ also performed family-wise error correction using cluster-level thresholding in each group map, and calculated the number of overlapping voxels passing correction. However, cluster-level correction allows for cluster-level, but not voxel-level inference. That is, the cluster is the unit of significance rather than the voxels within the cluster. Noting the number of overlapping voxels therefore does not capture the essence of whether a cluster has replicated or not. Therefore, I modified the procedure to determine the number of overlapping clusters rather than voxels. A cluster was deemed to have replicated if at least half of the voxels of that cluster were present in the replicate. Half is an arbitrary number intended to safeguard against trivial overlap. Finally, Turner et al^5^ examined peak overlap determined by whether the peak of a given cluster was also significant in the replicate. This is likely to be an important practical metric of replicability given that replication attempts will often examine a small radius around the peak of a previous report.

As in Turner et al^5^ each Jaccard overlap was performed at both a conservative threshold (depicted in the main text) and liberal threshold (depicted in the supplemental material). The liberal/conservative thresholds were as follows: voxel-level: p < 0.00025/0.00000025; cluster-level: p < 0.05 height, 1019 voxel extent/p < 0.01 height, 300 voxel extent, each achieving alpha < 0.01 according to 3dClustSim in AFNI. Interestingly, although it has been reported that liberal cluster-forming thresholds have inflated false positives^13^, which would be expected to harm replicability, replicability measures improved at the more liberal thresholds, which was also observed in Turner et al^5^ to some extent.

To quantify whether short or long scanning durations per task are the norm for the basic science domain from which the observed study is drawn, I searched PubMed for papers published since the start of 2015 using the terms “fMRI AND (cognitive control OR working memory)”. I excluded studies of special populations (e.g. patients, children) and interventional studies (e.g. drug, training) to focus on basic mechanistic research. The duration that each task was scanned was estimated from the reports. Functional localizer tasks producing regions-of-interest for a main task were excluded. The durations of the 244 resulting tasks are summarized in Supplemental Figure 3. The database is included at https://osf.io/b7y9n.

---

**Supplemental Figure 1.**
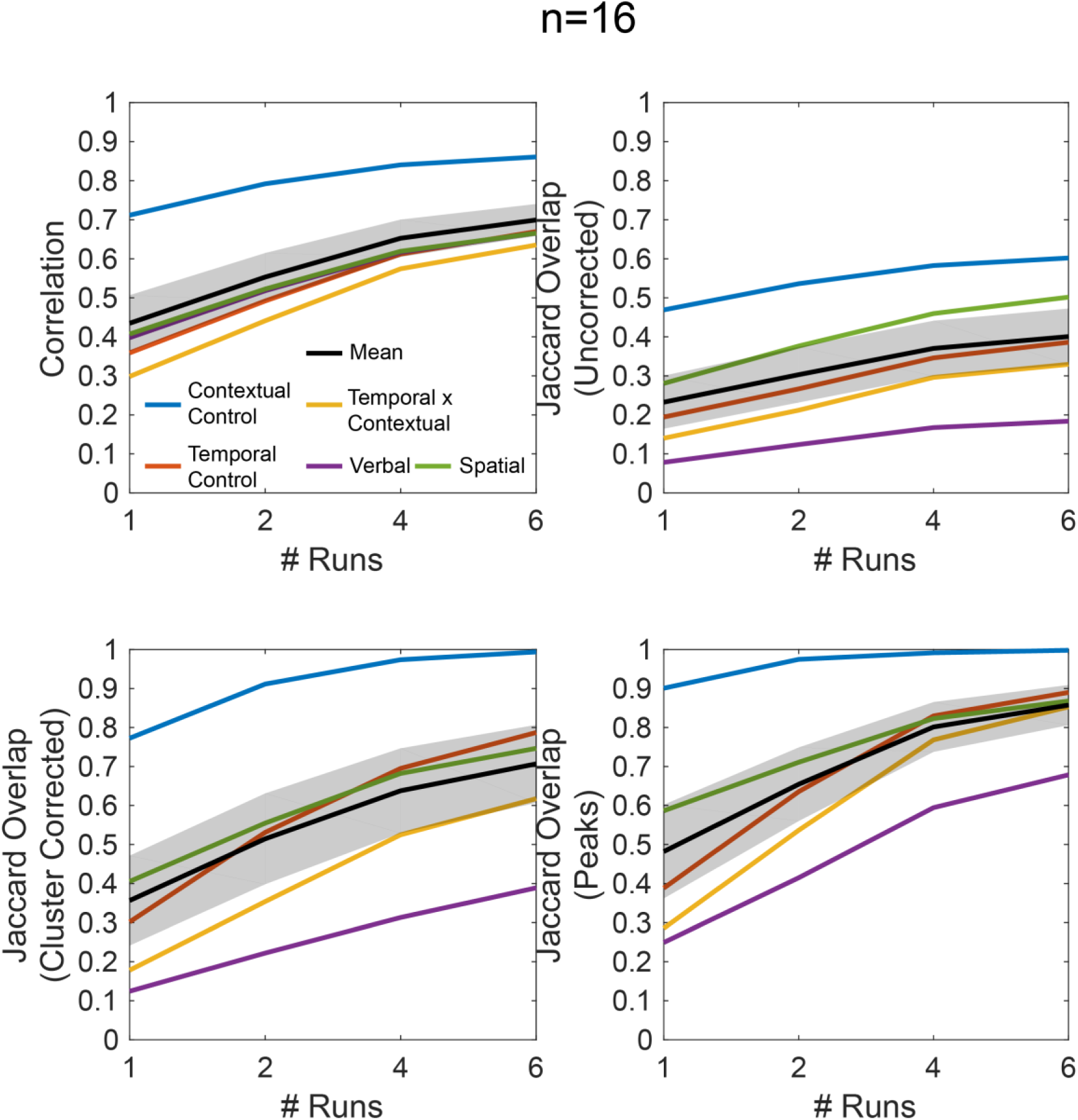
Details are identical to Figure 1, but Jaccard Overlap was computed using liberal threshold comparable to those reported in Turner et al^5^.

**Supplemental Figure 2.**
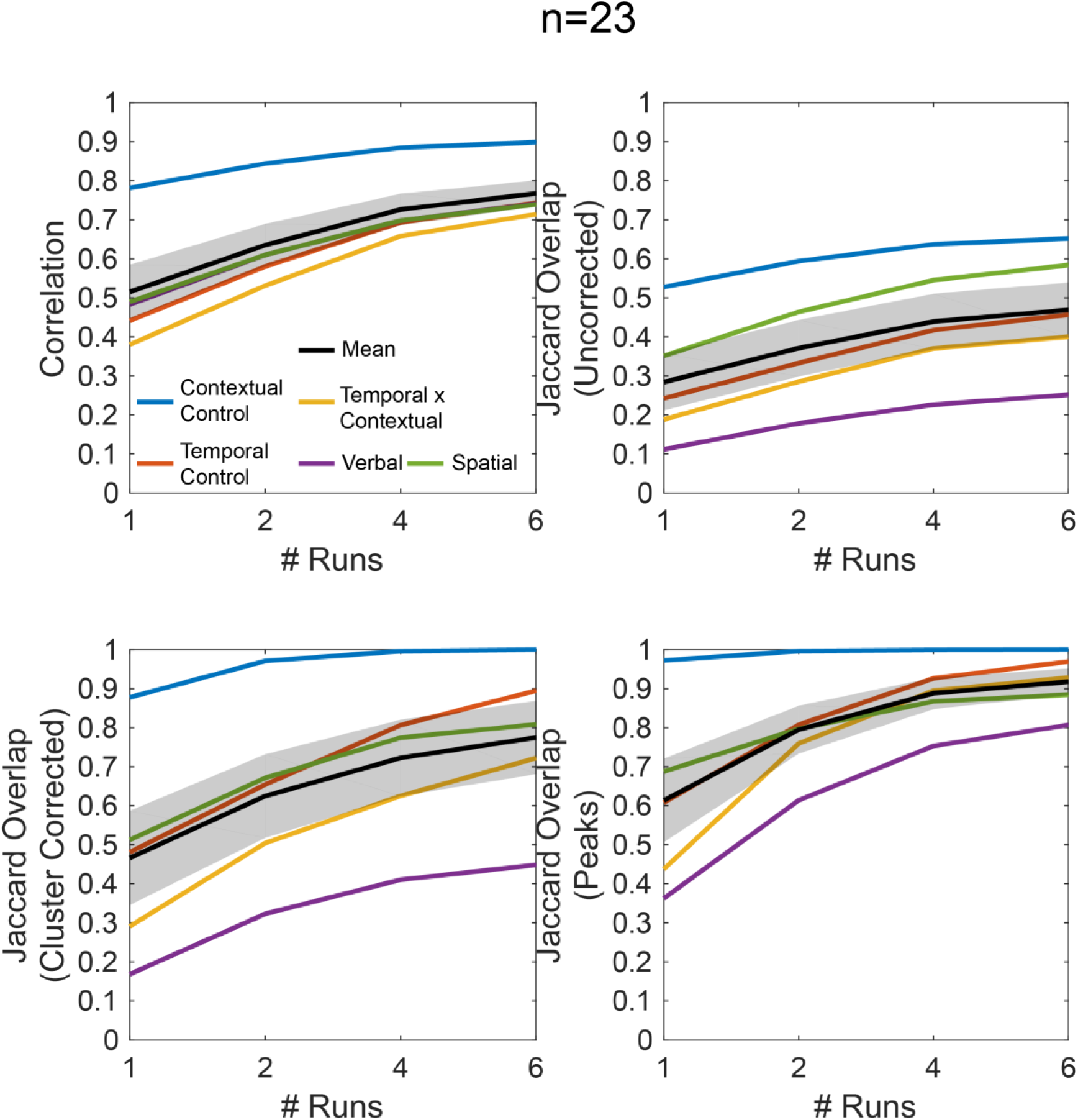
Replicability estimates at n=23 with liberal thresholding. Other details match Supplemental Figure 1.

**Supplemental Figure 3.**
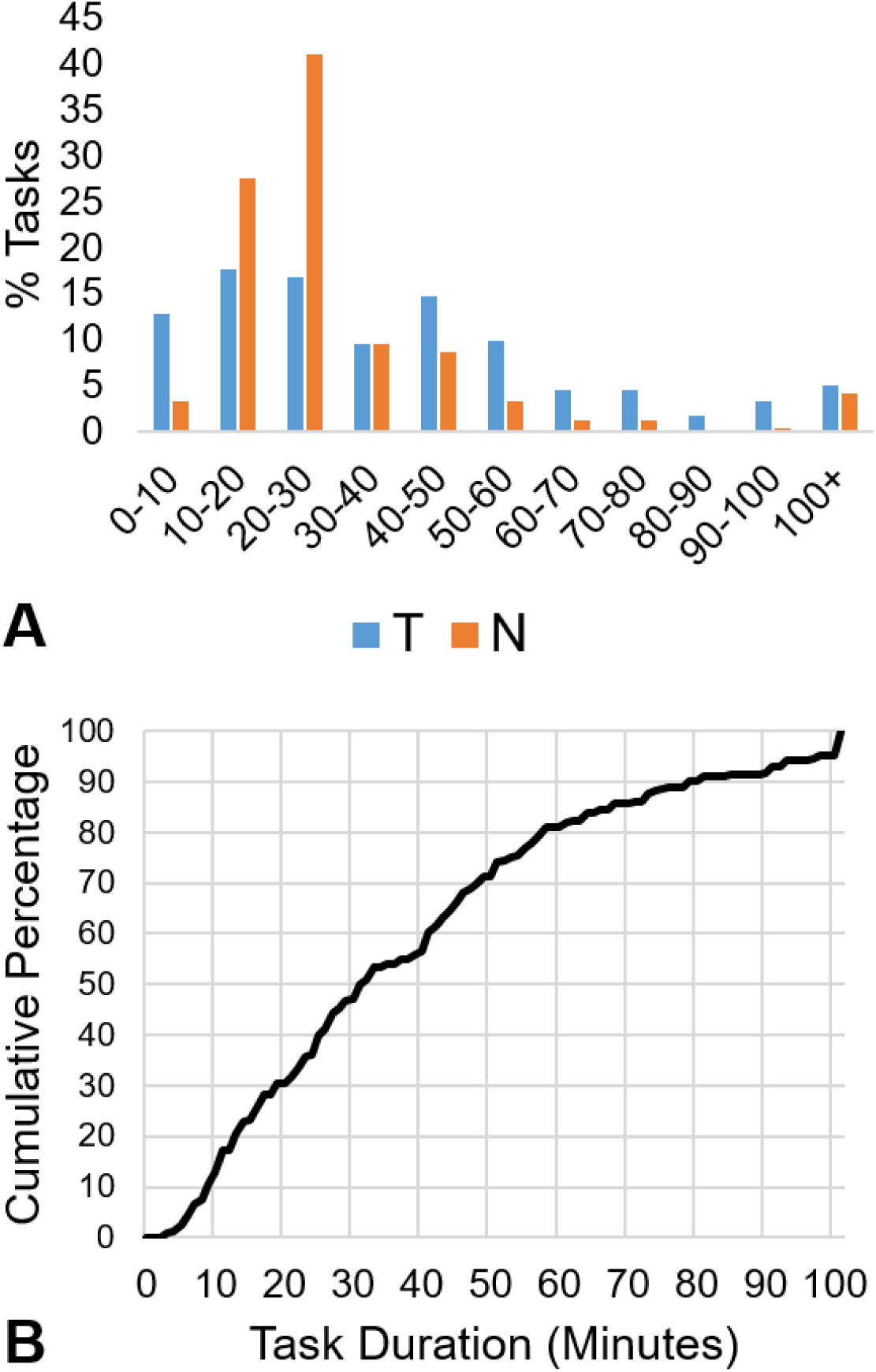
Estimates of scanning durations and sample sizes for basic science research in the domains of cognitive control and working memory since 2015. A) Histograms of task durations in minutes (T) and sample size (N). Sample sizes tend to be modest (N=20-30), but task durations less than ten minutes were uncommon (<15%). Mean (standard deviation) of task duration was 39.86 (33.20) minutes, and sample size was 31.73 (34.88) participants. B) Cumulative percentage of task durations.

